# Resource diversity structures aquatic bacterial communities

**DOI:** 10.1101/387803

**Authors:** Mario E. Muscarella, Claudia M. Boot, Corey D. Broeckling, Jay T. Lennon

**Affiliations:** Department of Plant Biology, University of Illinois, Urbana-Champaign 61801, USA; Department of Biology, Indiana University, Bloomington, Indiana 47405, USA; Natural Resource Ecology Laboratory, Colorado State University, Fort Collins, Colorado, USA; Department of Chemistry, Colorado State University, Fort Collins, Colorado, USA; Proteomics and Metabolomics Facility, Colorado State University, Fort Collins, Colorado, USA

**Keywords:** DOM Diversity, Resource Heterogeneity, Bacteria-DOM Interactions, Resource Specialization, Community Evenness, Microbial Generalists, Ecosystem Metabolomics

## Abstract

Microbial diversity is strongly affected by the bottom-up effects of resource availability. However, because resource pools often exist as heterogeneous mixtures of distinct molecules, resource heterogeneity may also affect community diversity. To test this hypothesis, we surveyed bacterial communities in lakes that varied in resource concentration. In addition, we characterized resource heterogeneity in these lakes using an ecosystem metabolomics approach. Overall, resource concentration and resource heterogeneity affected bacterial resource-diversity relationships. We found strong relationships between bacterial alpha-diversity (richness and evenness) and resource concentration and richness, but richness and evenness responded in different ways. Likewise, we found associations between the composition of the bacterial community and both resource concentration and composition, but the relationship with resource composition was stronger. Last, in the surveyed communities the presence of resource generalists may have reduced the effect of resource heterogeneity on community composition. These results have implications for understanding the interactions between bacteria and organic matter and suggest that changes in organic matter composition may alter the structure and function of bacterial communities.

## INTRODUCTION

Resource availability is a bottom-up control that has strong effects on the diversity of consumer communities. Theory suggests that resource enrichment promotes diversity and food-web complexity (Rosenzweig, 1971; Hairston and Hairston, 1993; Abrams and Roth, 1994; Polis and Strong, 1996; Worm *et al.*, 2002), and empirical studies have shown that, in the absence of top-down control, ecosystems with higher resource concentrations support more diverse and productive communities (Leibold *et al.*, 1997; Leibold, 1999; Hulot *et al.*, 2000; Waldrop *et al.*, 2006). However, the relationship between resources and diversity can be complex (Mittelbach *et al.*, 2001; Tilman *et al.*, 1982). For example, while diversity often increases linearly with resource concentration (Stevens and Carson, 2002), it can also exhibit more complex, non-linear relationships where diversity peaks at intermediate concentrations (Leibold, 1999). Such responses have been attributed to a range of processes including variation in competitive ability among consumers (Leibold, 1999), shared limitations across species (Stevens and Carson, 2002), and trophic interactions (Holt *et al.*, 1994; Carpenter *et al.*, 2001).

Another feature that may influence resource-diversity relationships is the heterogeneity of the resource pool. Resources are often considered as homogenous pools, but many resources exist as heterogeneous mixtures of multiple forms (Ashton *et al.*, 2010; Schoener, 1974; Turner, 2008). Resource heterogeneity has the potential to promote consumer diversity via niche partitioning (Schoener, 1974; Finke and Snyder, 2008). For example, plants have been shown to partition different forms of nitrogen (e.g., NH_4_, NO_3_, organic N) in ways that may promote species coexistence (McKane *et al.*, 2002; Schimel and Bennett, 2004; Andersen and Turner, 2013). Likewise, different phosphorus resources (e.g., phosphate vs. phytic acid) can alter the diversity and function of aquatic bacterial communities (Muscarella *et al.*, 2014), taxa in microbial biocrusts have non-overlapping resource preferences (Baran *et al.*, 2015), and phytoplankton are capable of partitioning the light spectrum in ways that allow for species coexistence (Stomp *et al.*, 2004).

The effects of resource heterogeneity may depend on the degree to which communities are comprised of generalist or specialists. If communities are made up primarily of resource generalists, then the total concentration of a resource should have a stronger influence on diversity because species do not differ in their response to different resources (Stevens and Carson, 2002). In contrast, if communities are made up of resource specialists, then resource heterogeneity may promote consumer diversity by providing unique resource niches for consumers to partition (Glasser, 1984; Levine and HilleRisLambers, 2009). Together, resource heterogeneity and resource acquisition strategy (i.e., generalists versus specialist) may help resolve unexplained variation in resource-diversity relationships.

For heterotrophic organisms, an important resource used for growth and physiological maintenance is organic matter. Organic matter is heterogeneous and consists of molecules that differ in chemical structure, origin, and age (Stevenson, 1994). In aquatic ecosystems, dissolved organic matter (DOM) is often classified based on origin (autochthonous vs. allochthonous) and bioavailability (labile vs. recalcitrant). DOM can also be characterized based on its optical properties (Fellman *et al.*, 2009; Weishaar *et al.*, 2003) and functional groups (e.g., humic acids) (Croué, 2004). However, these characterizations may not adequately describe DOM composition because other chemical features, including molecular weight, oxidation state, stoichiometry, and chemical structure, can influence the metabolism of organisms that consume DOM, (Cory and McKnight, 2005; Cherif and Loreau, 2007; Lennon and Pfaff, 2005; Berggren *et al.*, 2010). However, recent technological advances have made it possible to more thoroughly characterize DOM diversity at the molecular level (Moran *et al.*, 2016; Petras *et al.*, 2017; Broeckling *et al.*, 2008). Therefore, there is now the opportunity to understand the linkages between DOM heterogeneity and consumer diversity and characterize resource-diversity relationships (Töpper *et al.*, 2012; Alonso-Sáez and Gasol, 2007; Gómez-Consarnau *et al.*, 2012; Osterholz *et al.*, 2018).

In this study, we measured aquatic microbial communities and DOM chemistry to understand how resource heterogeneity contributes to resource-diversity relationships. We measured bulk resource concentration measurements and used high-resolution mass spectrometry to quantify resource heterogeneity. We also characterized aquatic bacterial community diversity using 16S rRNA sequencing. Furthermore, we used species-resource co-occurrence to test the hypothesis that communities dominated by specialists would respond stronger to resource heterogeneity than to resource availability. Our results support the view that resource heterogeneity promotes bacterial community diversity, but the contribution of DOM resource heterogeneity may be dampened when DOM generalists dominate bacterial communities.

## METHODS

### Study System and Sampling

The Huron Mountains nature preserve is a 5300 ha tract of private land in the upper peninsula of Michigan, USA. The area is part of the Superior Bedrock Uplands region (Schaetzl *et al.*, 2013). The surrounding forests are primarily old-growth hemlock-northern hardwoods (Woods, 2000), and the inland water bodies are part of the Pine River Watershed, which drains into Lake Superior. Using a van Dorn sampler, we obtained surface water samples (0.5 m) from 10 lakes in the Huron Mountains during July 2011 (Fig. S1, Table 1). In addition, we measured dissolved oxygen concentrations, temperature, pH, and conductivity at the time of sampling using Quanta Hydrolab water quality sonde, and we measured chlorophyll *a* concentration in the lab after cold ethanol extraction of 0.7 μm-filtered (Whatman GF/F) water samples using a Turner Biosystems Fluorometer (Table 1).

**Table 1.**
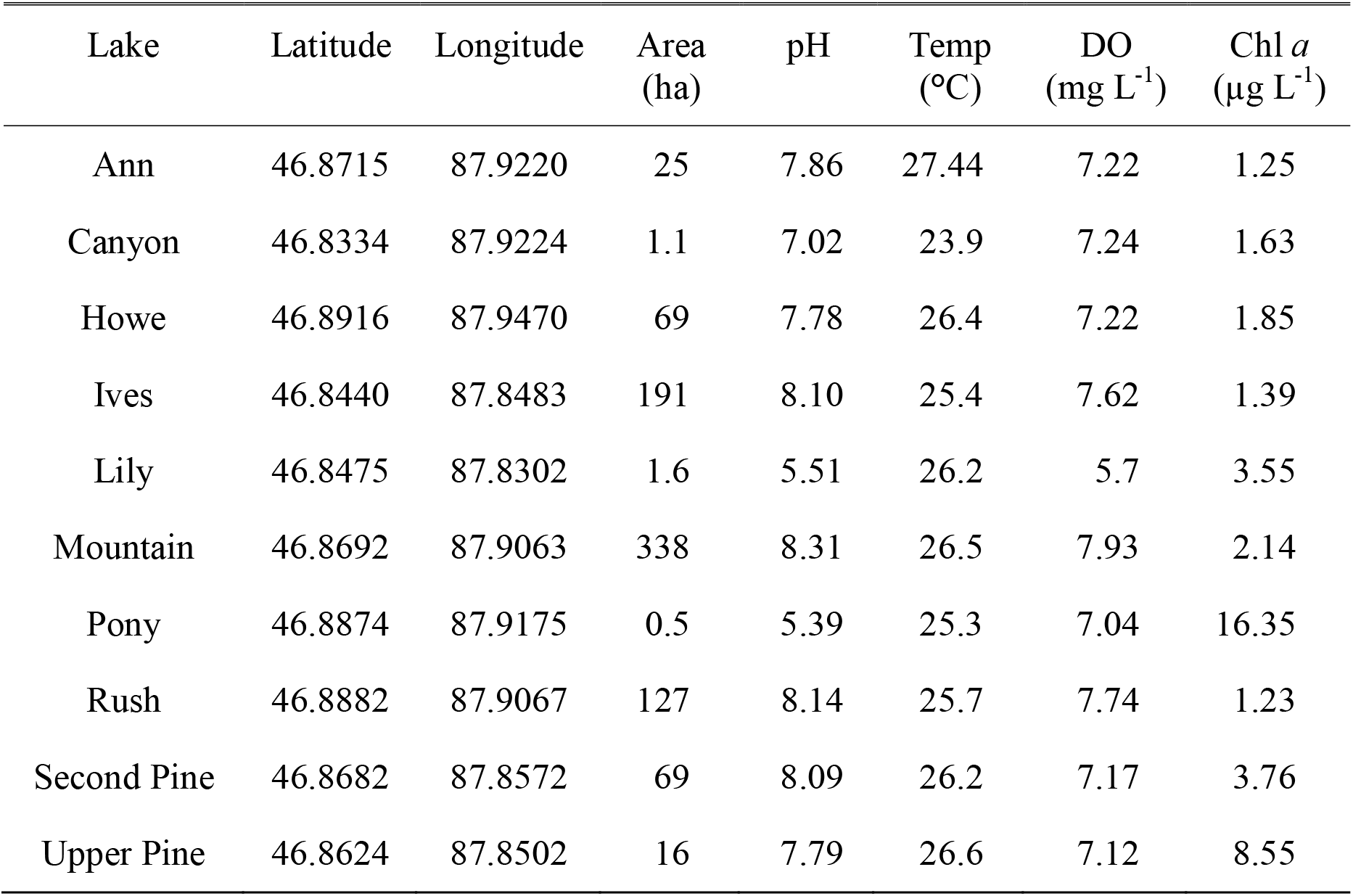
Lake Properties and Chemistry – Latitude, longitude and surface area (hectares), pH, temperature (Temp.), dissolved oxygen concentration (DO), chlorophyll *a* concentration (Chl *a*), TN: total nitrogen, TP: total phosphorus, DOC: dissolved organic carbon.

**Table 1 Cont.** Lake Chemistry. TN: total nitrogen, TP: total phosphorus, DOC: dissolved organic carbon.

| Lake | DOC (mg C L<sup>-1</sup>) | TP (µg P L<sup>-1</sup>) | TN (mg N L<sup>-1</sup>) |
| --- | --- | --- | --- |
| Ann | 5.97 | 7.27 | 0.43 |
| Canyon | 7.23 | 2.64 | 0.38 |
| Howe | 7.04 | 5.21 | 0.57 |
| Ives | 6.91 | 9.15 | 0.38 |
| Lily | 14.35 | 11.55 | 0.93 |
| Mountain | 5.27 | 5.87 | 0.34 |
| Pony | 28.99 | 17.04 | 1.86 |
| Rush | 4.22 | 3.84 | 0.41 |
| Second Pine | 6.26 | 12.92 | 0.44 |
| Upper Pine | 7.84 | 11.21 | 0.57 |

### Resource Concentrations

With the water samples, we measured the concentrations of dissolved organic carbon (DOC), total nitrogen (TN) and total phosphorus (TP). We measured DOC concentrations by oxidation and non-dispersive infrared detection on 0.7 μm-filtered (Whatman, GF/F) samples using a Shimadzu TOC-V carbon analyzer. We measured TN on unfiltered samples using a Lachat FIA 8500 auto-analyzer (Hach, Loveland CO) after ammonium peroxydisulfate/sulfuric acid digestion (Lachat, 2005). We measured TP on unfiltered samples spectrophotometrically using the ammonium molybdate method and oxidation by persulfate digestion (Wetzel and Likens, 2000).

### Resource Heterogeneity

To estimate resource heterogeneity, we characterized the composition of dissolved organic matter (DOM) for each lake using ecosystem metabolomics. We extracted DOM from each sample using solid phase extraction (SPE) (Dittmar *et al.*, 2008). Briefly, we acidified 1 L of 0.7 μm-filtered (Whatman, GF/F) water to pH 3.0 with 4N HCl. We then passed the water sample through an SPE cartridge (Discovery-18, Supelco, Bellefonte PA) at a flow rate ≤ 5 mL min^-1^ using vacuum pressure. Columns were pre-conditioned using 6 mL 100% methanol followed by 6 mL pH 3.0 ultra-pure H_2_O. We filtered the sample until no sample remained or until the cartridge became clogged (recording the final volume filtered) and dried the filter with N_2_ gas for 5 minutes. We eluted the DOM from the column using 100 % methanol and evaporated the methanol at 25 °C using vacuum centrifugation. A consistent amount of purified DOM was then separated on Waters Acquity ultra-performance liquid chromatography T3 column (1.8 μM, 1.0 x 100 mm) using water with a 0.1% formic acid-acetonitrile gradient and analyzed using negative electrospray ionization with quadrupole time of flight mass spectrometry (Q-TOF MS; Waters G2 Q-TOF) and indiscriminate tandem MS (idMS/MS) at the Colorado State University Proteomics and Metabolomics Facility. Q-TOF MS provides high resolution, accurate mass quantification and idMS/MS provides high collision energy fragmentation without precursor ion selection acquired concurrently with low-collision energy MS data. For each sample, raw data files were converted to .cdf format, and a matrix of molecular features as defined by retention time and ion mass (*m/z*) was generated using the XCMS package in R (Smith *et al.*, 2016) for feature detection and alignment. Raw peak areas were normalized to total ion signal, and the mean area of the chromatographic peak was calculated from duplicate injections. Features were grouped based on an in-house clustering tool, RAMClustR, which groups features into spectra based co-elution and covariance across the full dataset, whereby spectra are used to determine the identity of observed compounds in the experiment (Broeckling *et al.*, 2014). We used field-prepared ultrapure water as controls and subtracted control peaks from sample peak heights. We multiplied control peaks by 1.1 to provide conservative blank subtraction. A subset of the clustered dataset was referenced to the NISTv14 tandem (MS/MS) library, which contains 193,119 spectra of 43,912 precursor ions from 8,531 chemical compounds, and also screened for matches in Metlin for putative compound identification. Retention time was used as a proxy for polarity, and the heaviest ion (*m/z*) in the clustered spectra was used as a proxy for molecular weight. We recognize that the heaviest ion may not be representative of the molecular weight in all cases due to, for example, the formation of dimers and potential in-source fragmentation, however, electrospray is the gentlest type of ionization and is often related to the mass of the analytes. We define “DOM components” as the chemical features identified in DOM samples by ecosystem metabolomics.

### Microbial Composition

We used an RNA based approach to characterize bacterial community composition by sequencing the 16S rRNA gene transcript. We extracted total nucleic acids using the MoBio Power Water RNA extraction kit (Carlsbad, CA). Nucleic acid extracts were cleaned via ethanol precipitation and RNA extracts were treated with DNase I (Invitrogen) to degrade residual DNA. We synthesized cDNA via the SuperScript III First Strand Synthesis Kit using random hexamer primers (Invitrogen). Once cDNA samples were cleaned and quantified, we amplified the 16S rRNA gene transcript (cDNA) using barcoded primers (515F and 806R) designed to work with the Illumina MiSeq platform (Caporaso *et al.*, 2012). We purified the sequence libraries using the AMPure XP purification kit, quantified using the QuantIt PicoGreen kit (Invitrogen), and pooled libraries at equal molar ratios (final concentration: 20 ng per). After pooling, we sequenced the libraries on the Illumina MiSeq platform using 250 × 250 bp paired end reads (Illumina Reagent Kit v2) at the Indiana University Center for Genomics and Bioinformatics Sequencing Facility. Paired-end raw 16S rRNA sequence reads were assembled into contigs and filtered based on quality score, length, and ambiguous base calls. After filtering, we aligned our sequences to the Silva Database (version 123). Chimeric sequences were detected and removed using the VSEARCH algorithm (Rognes *et al.*, 2016). We then created operational taxonomic units OTUs by first splitting the sequences based on taxonomic class (using the RDP taxonomy) and the binning sequences in OTUs based on 97% sequence similarity. All initial sequence processing was completed using the software package mothur (version 1.40.5; Schloss *et al.*, 2009).

### Resource Heterogeneity and Community Diversity

First, we tested the hypothesis that resource heterogeneity affects bacterial community alpha diversity. We used linear models to determine if higher resource concentrations or more types of DOM resources (i.e., resource richness) would affect the richness and evenness of bacterial communities. We transformed (Box-Cox), centered, and scaled (i.e., divided by standard deviation) resource concentration and species richness to meet model assumptions of equal variance and normality (Neter *et al.*, 1996). We subsampled bacterial communities using rarefication to correct for differences in sample size due to sequencing depth (Hughes and Hellmann, 2005; James and Rathbun, 1981). We rarefied communities and calculated species richness as the number of OTUs observed and species evenness using Simpson’s evenness (Smith and Wilson, 1996). We used the Box-Cox-transformed DOC concentration as the measure of resource concentration and we calculated resource richness as the number of distinct DOM peaks observed in each sample.

Next, we tested the hypothesis that resource heterogeneity affects community beta diversity by comparing resource concentrations and DOM composition to bacterial community composition. We used distance-based redundancy analysis (dbRDA; Legendre and Anderson, 1999) to test for relationships between: 1) resource concentration and community composition and 2) resource composition and community composition. dbRDA is a multivariate linear model technique that uses quantitative factors explaining differences in multivariate community composition data. We used the Box-Cox-transformed DOC concentration as the measure of resource concentration. To use DOM composition as a predictor in our dbRDA model, we used principal coordinates analysis (PCoA), based on relative abundances and Bray-Curtis dissimilarity, to decompose DOM composition into orthogonal linear components (Legendre and Legendre, 2012). To represent the DOM composition, we used the DOM PCoA axis scores for each sample. As the response in the dbRDA model, we relativized OTU abundances and used Bray-Curtis distances to compare community composition across samples. Significance tests of our dbRDA model were conducted based on 10,000 permutations. All calculations were done in the R statistical environment (R Core Team, 2012) using the ‘vegan’ package (Oksanen *et al.*, 2013).

### Consumer-Resource Specialization

To test the hypothesis that the response to resource heterogeneity depends on whether communities were dominated by generalists or specialists, we used consumer-resource co-occurrence to define generalists and specialists. We defined resource generalists and specialists based on co-occurrence analysis, which was performed using Spearman’s rank correlations between DOM components and bacterial OTUs. We used the relative abundances of DOM components and the relative transcript abundances of bacterial OTUs. We inferred interactions based on correlations with coefficients > |0.7| (Williams *et al.*, 2014), and we tested for significance using a permutation test based on randomizations with the independent-swap algorithm (Gotelli, 2000). We defined resource generalists as those taxa with four or more significant negative resource interactions. We used the negative interaction as a proxy for potential resource consumption. To understand the spatial extent of individual taxa, we defined cosmopolitan taxa as those found in ≥ 90 % of the sampled lakes and we determined how many resource generalists were also cosmopolitan taxa. All calculations were done in the R statistical environment.

## RESULTS

### Resource Composition and Heterogeneity

The lakes sampled captured a range of bulk resource concentrations (Table 1), and many bulk resource concentrations were correlated. For example, the concentrations of dissolved organic carbon (DOC) and total nitrogen (TN) were highly correlated (rho = 0.97, *p* < 0.001, Fig. S2). Using ecosystem metabolomics, we characterized the dissolved organic matter (DOM) pool and detected 712 compounds across the sites. We refer to these molecules as DOM components. Based on the relative abundances of DOM components, sites were on average 37 % dissimilar in DOM composition. Using principal coordinates analysis (PCoA), we could explain 71 % of the variation in DOM composition across sites using three dimensions (Fig. 1). The variation in DOM composition was significantly related to DOC (r^2^ = 0.68, *p* = 0.01), TN (r^2^ = 0.70, *p* = 0.01), Chl a (r^2^ = 0.69, *p =* 0.02), and pH (r^2^ = 0.58, *p* = 0.03), but there were no significant relationships with TP (r^2^ = 0.27, *p* = 0.34) or surface area (r^2^ = 0.30, *p* = 0.26). In addition, we found a negative relationship between the richness of DOM components and the concentration of DOC (p < 0.01). We used DOC to represent resource concentration and the DOM PCoA scores to represent DOM composition in further analyses. We identified influential DOM components as those correlated (rho > |0.70|) with variation in the DOM PCoA axes (Fig. S3). We identified 172 influential DOM components.

**Fig 1:**
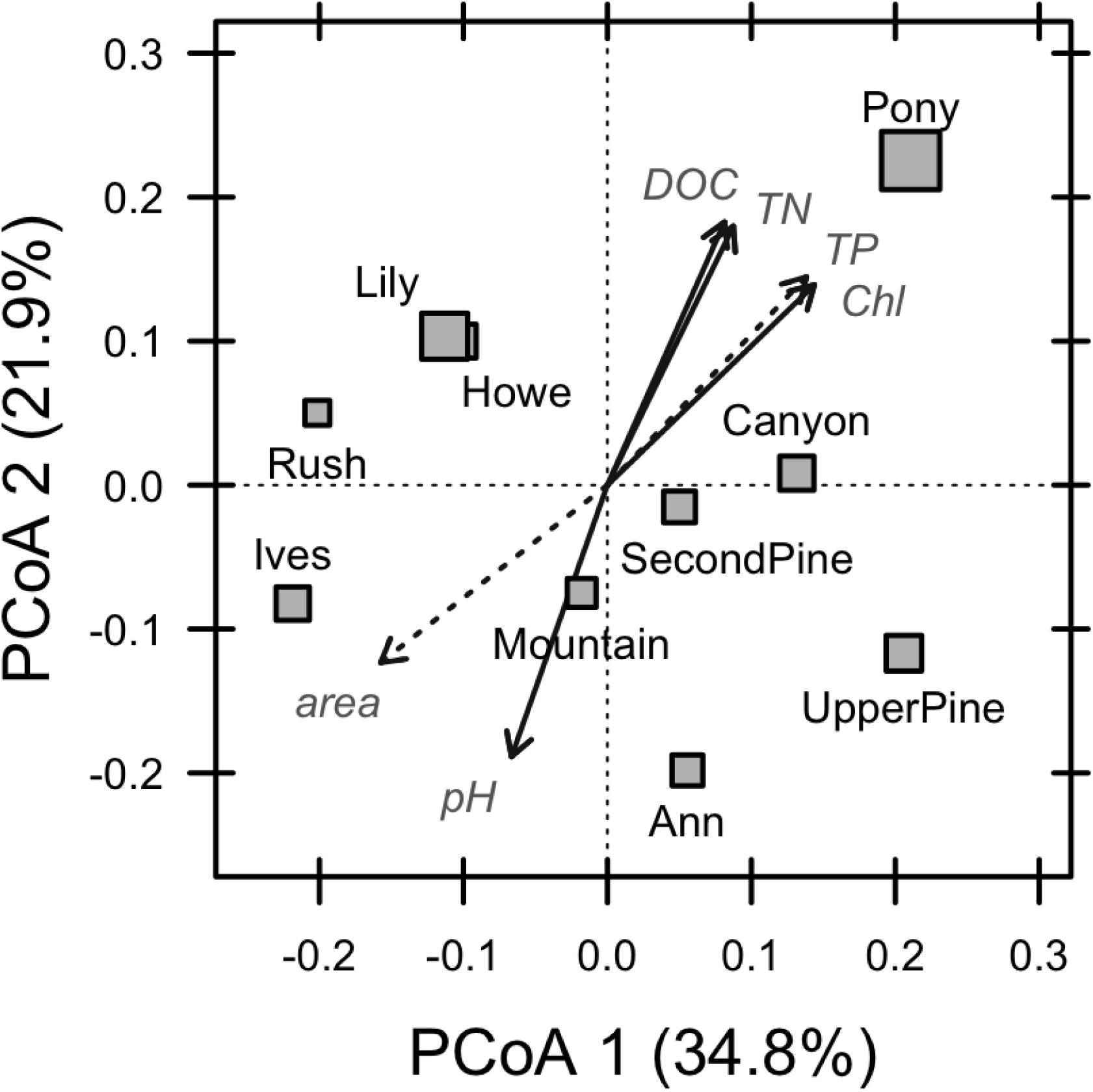
Principal coordinates analysis (PCoA) ordination of dissolved organic matter (DOM). The distances between symbols represent the dissimilarity between DOM in each lake. Using three axes, we can explain 71% of the variation in DOM composition. The third axis (not shown) captures 14% of the variation. Symbol sizes reflect variation in the concentration of dissolved organic carbon (DOC). Vectors represent the correlations between DOM composition and various physical and chemical attributes of each lake including: pH, area, DOC, total nitrogen (TN), total phosphorus (TP), and chlorophyll a (Chl).

### Community Composition and Resource-Diversity Relationships

Across the 10 lakes, we identified 5,085 bacterial operational taxonomic units (OTUs) based on 16S rRNA transcript sequencing. When rarified, lakes varied in taxonomic richness and evenness (Fig. 2). Using Bray-Curtis distances and relative transcript abundances, lakes were on average 62 % dissimilar to one another based on bacterial community composition.

**Fig 2:**
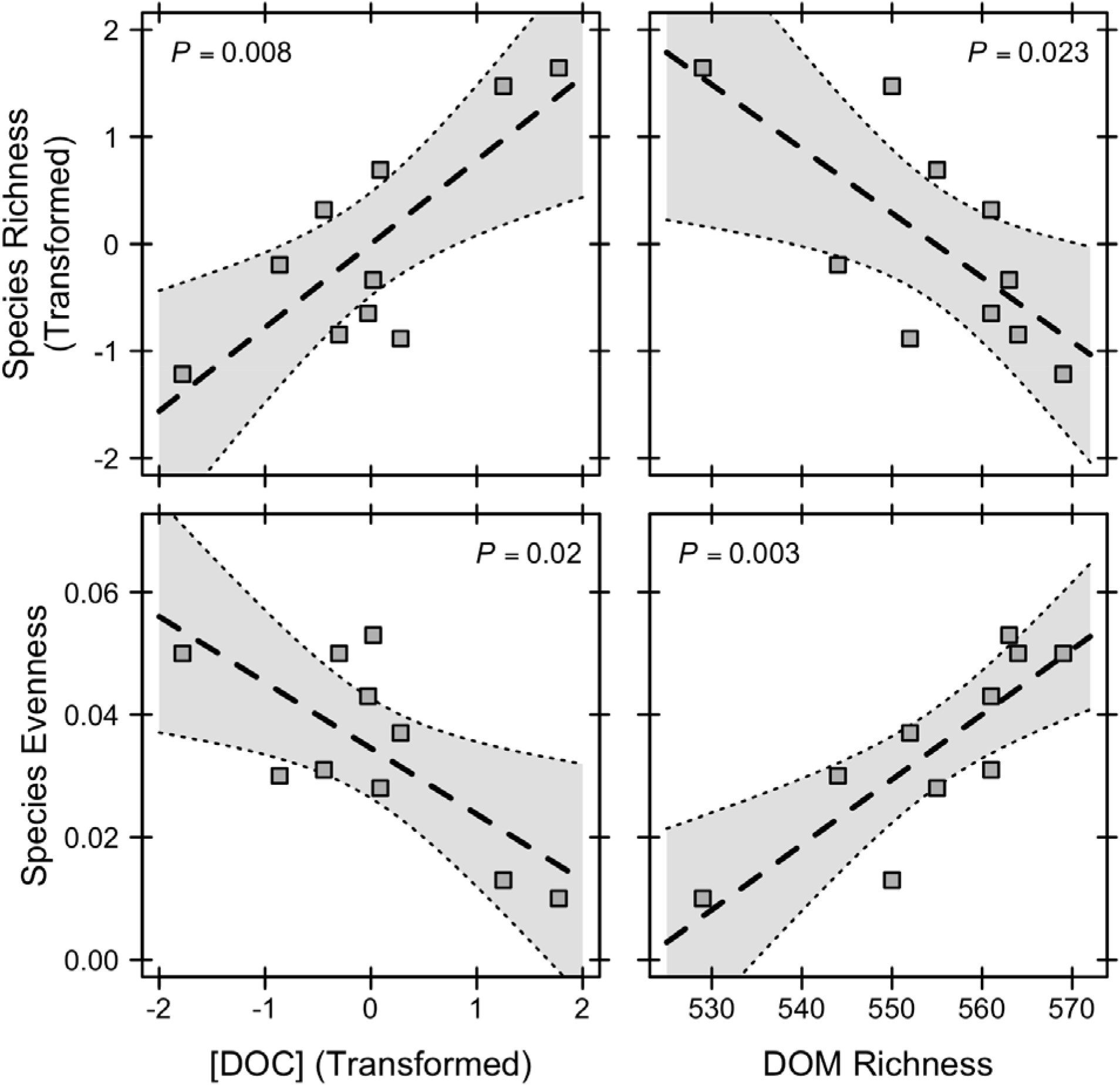
Bacterial community diversity relationships with resource concentration and resource richness. Resource (DOC) concentration and species richness have been Box-Cox transformed to meet model assumptions. There are significant positive relationships between species richness and resource concentration and between species evenness and dissolved organic matter (DOM) richness. There are significant negative relationships between species evenness and resource concentration and between species richness and DOM richness. Dashed line represents linear regression fit along with 95% confidence intervals.

First, we tested for relationships between resources and bacterial alpha-diversity. We used linear regression to test for resource-diversity relationships between bacterial community diversity (richness and evenness) and both resource concentration and DOM richness. As predicted, bacterial alpha-diversity was affected by resource concentration (Fig. 2). OTU richness was positively related to resource concentration (r^2^ = 0.66, *p* = 0.008) but negatively related to DOM richness (r^2^ = 0.50, *p* = 0.023). In contrast, OTU evenness was positively related to DOM richness (r^2^ = 0.67, *p* = 0.003) but negatively related to resource concentration (r^2^ = 0.49, *p* = 0.022).

Next, we tested for relationships between resources (concentrations and composition) and bacterial beta-diversity using distance-based redundancy analysis (dbRDA). Based on the dbRDA models, resource concentrations explained 28 % of the variation in bacterial community composition (*p* = 0.002), and DOM composition explained 45 % of the variation in bacterial community composition (p = 0.03, Fig. 3). However, when we partitioned the explained variation among the DOM PCoA axes, only DOM Axis 2 was significant (r^2^ = 0.70, *p* = 0.017). In addition, this DOM axis was correlated to variation along OTU PCoA Axis 1 (rho = 0.83, *p* = 0.002; Fig. 3). Last, we tested for relationships between resource concentration and DOM composition. We found a significant correlation between resource concentration and DOM Axis 2 (rho = 0.69, *p* = 0.03).

**Fig 3:**
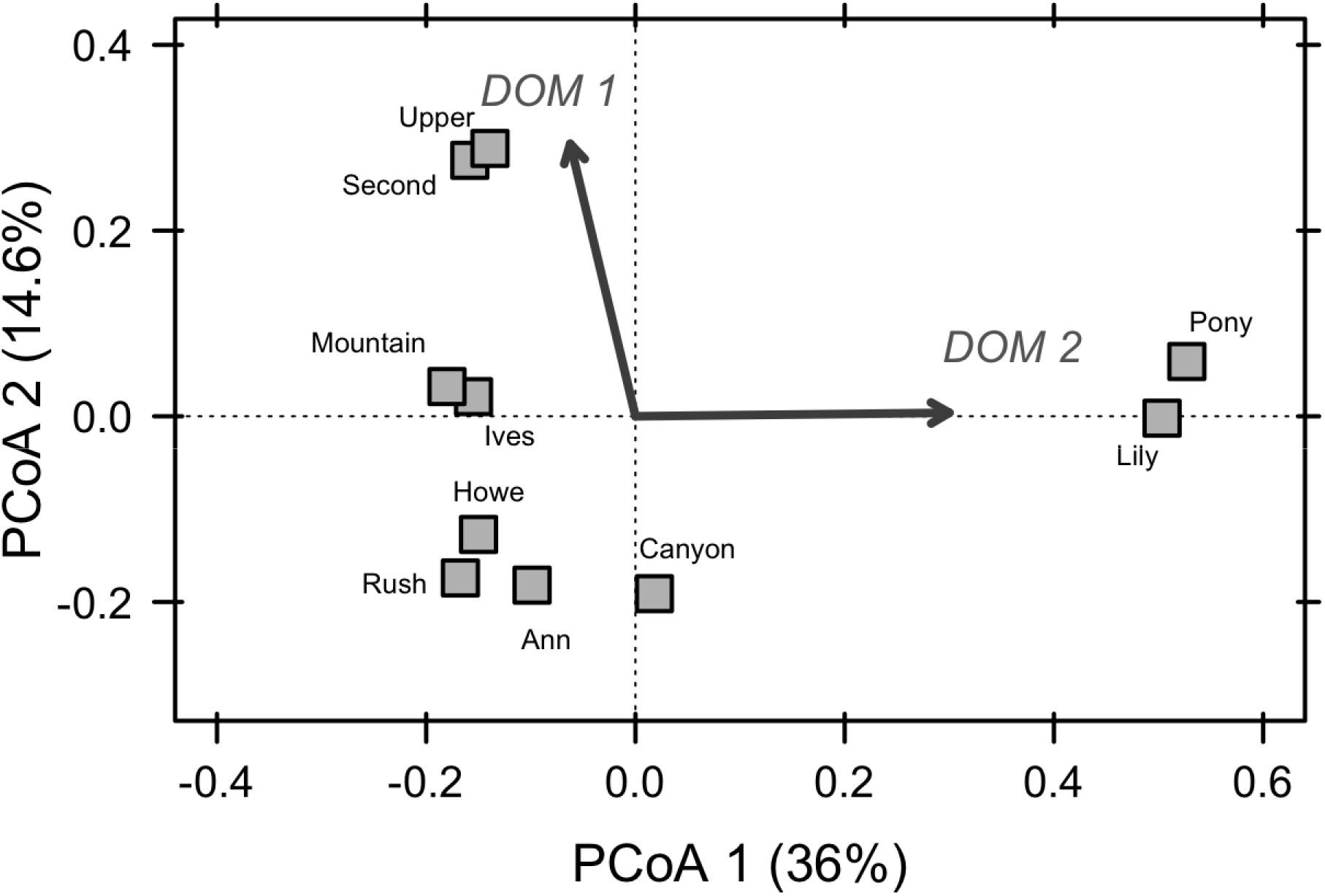
Principal coordinates analysis (PCoA) ordination of bacterial communities. Vectors represent the correlation between the dissolved organic matter (DOM) heterogeneity and the bacterial community composition. The two vectors are based on correlations between community composition and the site scores from the DOM PCoA axes one and two. We used distance-based redundancy analysis to test the relationship between DOM site scores and bacterial community composition.

### Consumer–Resource Specialization

Based on consumer-resource co-occurrence analysis and spatial occurrence, we classified generalist and cosmopolitan bacteria. We found that 1.3% of taxa (68 OTUs) were resource generalists, and 4.5 % (233 OTUs) were cosmopolitan taxa. Of the resource generalists, 74 % were also found to be cosmopolitan taxa. Proportionally, resource generalists and cosmopolitan taxa were substantial across all lakes (Fig. 4). For both groups, there was a significant negative relationship between relative abundance and resource concentration (Fig. 4). In addition, the proportion of resource generalists was related to DOM composition based on DOM Axis 2 (rho = 0.81, *p* = 0.004), but not DOM Axis 1 (rho = 0.08, *p* = 0.82). Taxonomically, both resource generalists and cosmopolitan taxa were diverse. For the resource generalists, the majority belonged to the classes Alphaproteobacteria (14) and Planctomycetacia (11), but Verrucomicrobiae (8) and Actinobacteria (7) were also common. At the family level, the resource generalists represented groups including Acetobacteraceae, Caulobacteraceae, Planctomycetaceae, Sphinomonadaceae, and Verrucomicrobiaceae. For the cosmopolitan taxa, the majority belonged to the classes Alphaproteobacteria (58) and Betaproteobacteria (50), but Gammaproteobacteria (14), Actinobacteria (21), Planctomycetacia (17), and Sphingobacteria (12) were also common. At the family level, the cosmopolitan taxa represent groups including Acetobacteraceaea, Alcaligenaceae, Bulkholderiaceae, Caulobacteraceae, Chitinophagaceae, Comomonadaceae, Flavobacteriaceae, Planctomycetaceae, Rhodobacteraceae, Spartobacteria, Sphinomonadaceae, and Verrucomicrobiaceae.

**Fig 4:**
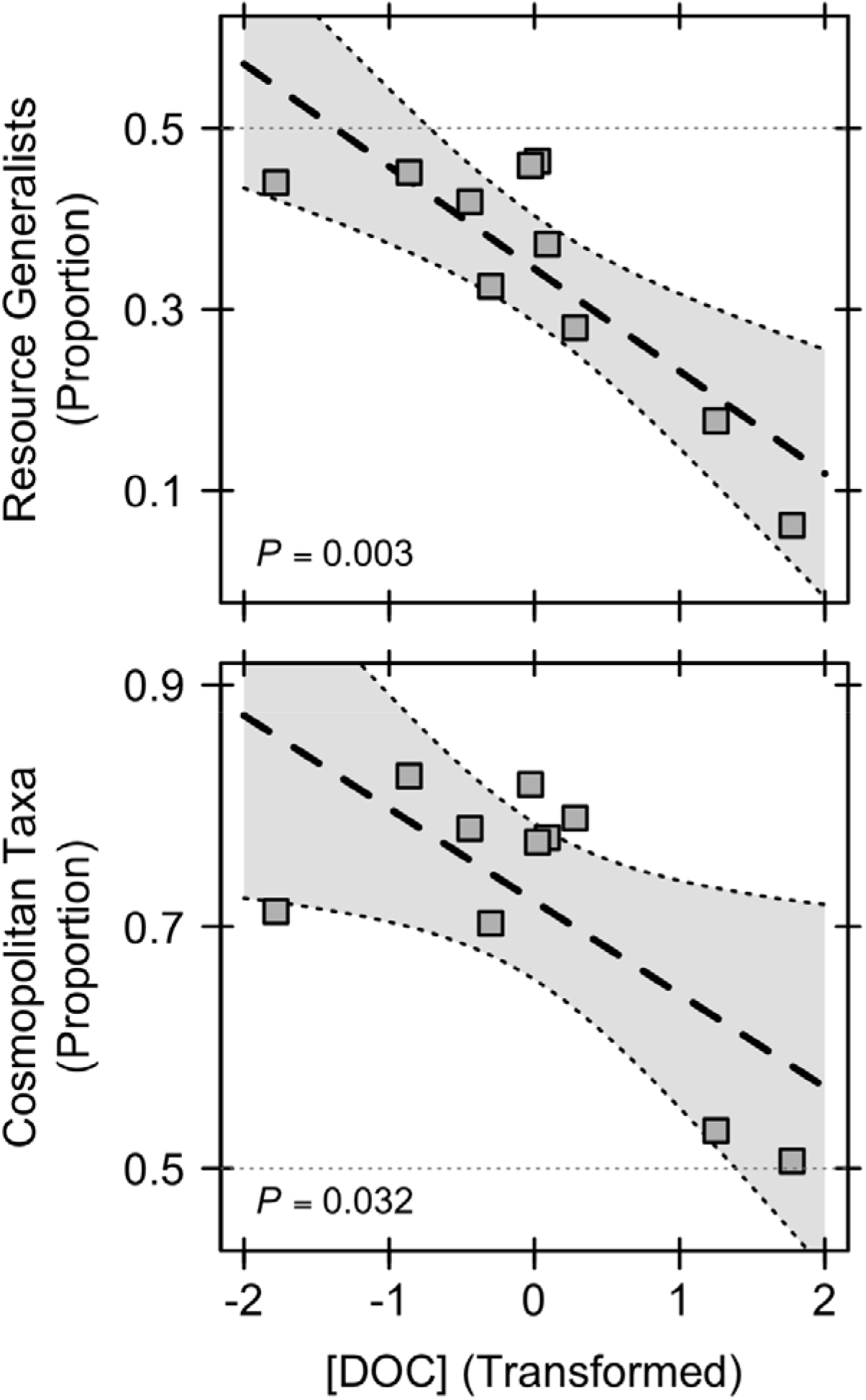
The proportion of generalists and cosmopolitan taxa in aquatic bacterial communities. We defined operational taxonomic units (OTUs) as generalists using consumer-resource co-occurrence (top) and as cosmopolitan based on spatial occurrence (top). We used OTU relative abundances to calculate the proportion in each community. For both, we used a linear model to determine if there was a relationship between the proportion of generalists and the concentration of dissolved organic carbon (DOC). For both, we found a significant negative relationship. Dashed line represents linear regression fit along with 95% confidence intervals. The light gray dotted line represents 50% of the community and is used as a reference.

## DISCUSSION

We found evidence that resource concentration and resource heterogeneity affect bacterial resource–diversity relationships. Our data suggest that there is a significant relationship between resources (concentration and richness) and bacterial community alpha diversity. Likewise, resource concentration and composition explained variation in bacterial community composition (beta-diversity), although to differing degrees. Last, DOM generalists were prevalent in the surveyed microbial communities and that there was a negative relationship between the proportion of generalists and the concentration and composition of DOM. Together, our results suggest that DOM resource heterogeneity affects aquatic microbial communities, and that DOM resources may influence aspects of community diversity (e.g., species evenness) and community composition. However, when generalists dominate communities, the effects may be limited potentially due to complex food-web interactions. Based on our findings, we argue that organic matter composition plays an important role in structuring aquatic microbial communities, and that changes in organic matter composition owing to land use modifications and changing terrestrial plant communities may alter the structure and function of aquatic bacterial communities.

### Resource Heterogeneity in Microbial Food Webs

Resource heterogeneity affected the diversity of aquatic bacterial communities. We found that DOM resources were heterogeneous across lakes — on average lakes were 37 % dissimilar in their DOM composition. As such, resource heterogeneity may help explain the variation in resource-diversity relationships along resource concentration gradients. We tested this hypothesis and found that while resource concentration explained 28 % of the variation, DOM resource composition explained 45 % of the variation in bacterial community composition across lakes. These findings suggest that different types of bacteria use and potentially specialize on different types of resources, which has been observed elsewhere. For example, it has been shown that some bacteria primarily use algal-derived resources (Sarmento and Gasol, 2012; Jaspers and Overmann, 2004) while others primarily use terrestrial-derived resources (Guillemette *et al.*, 2015; Roehm *et al.*, 2009). Therefore, lakes receiving different resource inputs may be expected to contain different bacterial communities. Thus, DOM resource heterogeneity is a potential mechanism to explain the diversity within and between bacterial communities.

Resource diversity (i.e., DOM richness) was positively correlated with OTU evenness, but negatively correlated with OTU richness (Fig. 2). Resource diversity is likely to influence OTU evenness because evenness, a measure of equitability among taxa, may reflect the frequency of species traits (Hillebrand *et al.*, 2008; Hill, 1973), such as enzymes needed to uptake and metabolize different DOM components. Furthermore, changes in evenness have been linked to altered species-interactions, coexistence, and ecosystem functions (Hillebrand *et al.*, 2008). If resources represented niches to be partitioned, resource diversity should promote species diversity because resource diversity provides unique niches to species to partition (Werner, 1977; Glasser, 1984; Schoener, 1974). Because we observed an increase in evenness but not in richness with greater resource diversity, our findings suggest that the increased evenness observed in communities represents changes in abundances but not the addition of new taxa. Furthermore, the change in evenness — an increase — suggests that the changes in abundance benefit intermediate rank taxa. Together, our results support the hypothesis that resource heterogeneity contributes to observed resource-diversity relationships. In addition, we propose that DOM resource heterogeneity may promote more diverse communities by increasing species equitability and benefiting taxa that comprise the middle ranks of the bacterial community - “The Microbial Middle Class”.

### Resource Substitutability

One possible explanation for why resource heterogeneity may only have weak effects in some habitats is that many resources are substitutable. Two resources are substitutable when either can each be used for growth and reproduction while the other is absent (Tilman, 1980). For example, some plants are able to grow using ammonium, nitrate, or even organic nitrogen as a source of nitrogen (Haynes and Goh, 1978; McKane *et al.*, 2002; Schimel and Bennett, 2004), and zooplankton such as *Daphnia* can use algae, cyanobacteria, and bacteria independently as food sources (Demott, 1998). Likewise, aquatic ecosystems contain numerous phosphorus resources but some have similar effects on the structure and function of aquatic microbial communities (Muscarella *et al.*, 2014).

In this study, we found numerous DOM components that appear to have similar consumer-resource co-occurrence patterns (Fig. S4). One explanation is that many DOM components are substitutable. At a chemical level, resources with the same core molecule can be substitutable. For example, vanillate and ferulate share an internal benzene structure and are used by the same metabolic pathways (Buchan *et al.*, 2000). In addition, extracellular enzymes often degrade aliphatic polymers of different lengths into identical monomers (Rojo, 2009). As such, many DOM components are likely substitutable. We used chemical databases to identify the resources captured by our mass spectrometry and group possible substitutable resource; however, we were unable to make positive identifications for many resource components in part due to the low representation of environmental samples in the available databases. In addition, we tested for patterns based on the polarity and molecular weight estimates but found no significant relationships. This does not mean relationships between DOM and bacterial composition are indescribable, but methods need to be developed to classify and group DOM components into meaningful categories based on functional and metabolic forms.

### Generalist Communities

Our results suggest that resource generalists may dominate many aquatic microbial communities, and thus may explain why the effect of resource heterogeneity on community composition is stronger is some lakes than others. Specifically, resource heterogeneity had a weak effect on the composition of bacteria in lakes that separate along OTU PCoA Axis 2 (Fig. 3). Across our lakes, we found a negative relationship between the abundance of generalists and the concentrations of resources. This relationship is also correlated with the second axis of the DOM PCoA, but not the first DOM PCoA axis which explains the majority of the DOM variation. One possibility is that the majority of DOM resources are substitutable. Alternatively, consumers could have multiple metabolic pathways for resource acquisition. For example, evidence from comparative genomics suggests that aquatic bacteria capable of using complex organic matter also have the potential to use numerous different resources, and may thus be generalists (Livermore *et al.*, 2013; Newton *et al.*, 2010; Lauro *et al.*, 2009). As such, we propose that resource generalists may be more common in aquatic ecosystems than previously thought (Mariadassou *et al.*, 2015).

It is often assumed that most bacteria are specialists. For example, multiple studies have identified taxa that specialize on particular resources (Hunt *et al.*, 2008; Mccarren *et al.*, 2010; Gómez-Consarnau *et al.*, 2012; Jaspers and Overmann, 2004; Bird, 2012). The ability to use multiple resources requires the production of extra enzymes and transporters; therefore, it is costly to use numerous resources (Johnson *et al.*, 2012). As such, specialists may be energetically favored in some environments. Likewise, numerous studies have indicated that habitat specialists (e.g., sediment and aquatic) dominate bacterial communities (Székely and Langenheder, 2013; Mariadassou *et al.*, 2015; Langenheder and Ragnarsson, 2007). However, our results suggest that generalists are common in the lakes surveyed (Fig. 4). These findings are supported by another study which found that resource generalists dominated coastal bacterial communities (Mou *et al.*, 2008). It should be noted though, that we found both resource generalists and specialists (Fig. S4) and therefore we are not suggesting that resource specialists do not contribute to resource-diversity relationships. Instead, we argue that generalist may limit the ability of resource heterogeneity to promote diversity when generalists are more dominant than specialists.

We do acknowledge, however, that the method used to characterize DOM has some limitations. First, the DOM extraction and detection may be biased towards some groups of molecules (Dittmar *et al.*, 2008). While we may have missed some important components of the DOM pool, we likely captured the complex terrestrial-derived organic matter that often dominates aquatic ecosystems (Wilkinson *et al.*, 2013). This DOM has been shown to be important for bacterial community structure and function (Lapierre *et al.*, 2013; Lennon and Pfaff, 2005; Muscarella *et al.*, 2016). However, these are not the only important components of the DOM pool, and we may have missed less complex labile molecules that can also affect bacterial communities (Sarmento and Gasol, 2012). However, many labile molecules would be consumed rapidly, and thus we may not have been able to detect them. Second, our consumer-resource interaction results are based on a single time point and therefore only suggest possible bacteria-resource interactions. We use these correlations to make inferences about the degree to which taxa are generalists. To make stronger inferences, we would need to conduct time-course experiments during resource fluctuations and perform experimental manipulations of DOM concentration and composition. Last, we assume that microbial communities are under local selection due to resource availability, but other factors such as dispersal, predation, and the physical environment can affect community composition. For example, high dispersal rates can overwhelm local selection due to mass effects (Leibold *et al.*, 2004) which is especially important in aquatic microbial communities that receive organisms from the neighboring terrestrial landscape (Ruiz-Gonzalez *et al.*, 2015; Crump *et al.*, 2012). Regardless, our results, and other genomic studies, suggest that resource generalist may dominate aquatic microbial communities, and this should be investigated further.

### Conclusions

Resource heterogeneity influenced the resource-diversity relationship and the contribution of heterogeneity can be greater than concentration; however, when resource generalists dominated communities the resource-diversity relationship was dampened. These findings do not mean that there are no specialists in bacterial communities, because we find evidence of resource specialist and others have found strong evidence for resource and habitat specialists (Székely and Langenheder, 2013; Mariadassou *et al.*, 2015; Langenheder and Ragnarsson, 2007; Bird, 2012; Muscarella *et al.*, 2016). These findings support the hypothesis that generalist taxa may limit the affect resource heterogeneity has on local communities; furthermore, we propose that consumer properties (i.e., generalist) and resource properties (i.e., availability) determine how strong communities respond to resource heterogeneity. In addition, in order to understand how bacterial communities will respond to environmental changes, such as changes in organic matter inputs due to changes in plant community distributions or global climate change, we need to consider which resources are substitutable and which resources will change in similar and predictive ways. In doing so, we will be able to understand how microbial communities will respond to alterations in the available resources.

## Acknowledgements

We acknowledge constructive feedback from AL Peralta and NI Wisnoski. We thank BK Lehmkuhl for technical support and W Thorpe for logistical support. This work was supported by the Huron Mountain Wildlife Foundation (MEM, JTL), the National Science Foundation DEB-1442246 (JTL) & DEB-1501164 (MEM, JTL), and US Army Research Office Grant W911NF-14-1-0411 (JTL). All code and data used in this study can be found in a public GitHub repository (https://www.github.com/LennonLab/ResourceHeterogeneity) and the NCBI SRA.

